# Multifractality in postural sway supports quiet eye training in aiming tasks: A study of golf putting

**DOI:** 10.1101/2020.08.19.258053

**Authors:** Noah Jacobson, Quinn Berleman-Paul, Madhur Mangalam, Damian G. Kelty-Stephen, Christopher Ralston

## Abstract

The ‘quiet eye’ (QE) approach to visually-guided aiming behavior invests fully in perceptual information’s potential to organize coordinated action. Sports psychologists refer to QE as the stillness of the eyes during aiming tasks and increasingly into self- and externally-paced tasks. Amidst the ‘noisy’ fluctuations of the athlete’s body, quiet eyes might leave fewer saccadic interruptions to the coupling between postural sway and optic flow. Postural sway exhibits fluctuations whose multifractal structure serves as a robust predictor of visual and haptic perceptual responses. Postural sway generates optic flow centered on an individual’s eye height. We predicted that perturbing the eye height by attaching wooden blocks below the feet would perturb the putting more so in QE-trained participants than participants trained technically. We also predicted that QE’s efficacy and responses to perturbation would depend on multifractality in postural sway. Specifically, we predicted that less multifractality would predict more adaptive responses to the perturbation and higher putting accuracy. Results showed that lower multifractality led to more accurate putts, and the perturbation of eye height led to less accurate putts, particularly for QE-trained participants. Models of radial error (i.e., the distance between the ball’s final position and the hole) indicated that lower estimates of multifractality due to nonlinearity coincided with a more adaptive response to the perturbation. These results suggest that reduced multifractality may act in a context-sensitive manner to restrain motoric degrees of freedom to achieve the task goal.

## 1. Introduction

Aiming our behaviors into the visible world requires an ongoing awareness of how our body fits in the world around it. The difficulty of sorting out this bodily fit finds its classic encoding in the ‘degrees of freedom’ problem: even if a fine-grained choreography of goal-directed movement that stipulates all individual motoric degrees of freedom were possible, this fine-grained choreography would probably be too rigid to adapt to the task or context (Bernstein, 1967). However, another perspective suggests that the abundance of degrees of freedom is not a problem but a ‘blessing’ (Latash, 2012). According to this view, there is no need to stipulate the infinitude of motoric variables at very fine grains, and on the contrary, task or intention organizes the movement system at the relatively coarse grain of the whole body and context (Latash, 2020). The infinitude thus becomes a guarantee of adaptive flexibility. This positive view of motor abundance appears in recent studies of aiming behaviors like the golf swing, whereby a clearly defined target in the context draws visual attention away from specific postures of the body, and indeed, experts with their eyes on the target show no single optimized mode of execution (Morrison et al., 2016). Hence, instead of requiring explicit choreography, the movement system’s organization may readily emerge from perceptual linkages to the surrounding context (Stoffregen, 1985, 1986; Stoffregen & Riccio, 1990).

### 1.1. Quiet eye vs. technical training

Supporting task performance by emphasizing perceptual linkages vs. explicit choreography is a clear theme of research on ‘quiet eye’ (QE). Sports psychologists refer to QE as the stillness of the eyes during aiming tasks, specifically the final gaze fixation to a single location or object in the task for a minimum of 100 ms within 3 degrees of visual angle immediately before the movement is initiated (Rienhoff et al., 2016). In the case of a golf swing, ‘technical training’ that communicates an explicit choreography of the arm and leg placement leads to less accuracy than a QE strategy. Expert golfers show more frequent and longer gaze fixations on specific aspects of the putting situation (e.g., golf ball, putting surface, target hole). Novices tend to fixate more introvertedly (e.g., on the clubhead during the backswing phase) and focus on the ball only after it has been hit with the club (Moore et al., 2012; Vickers, 1992, 1996; Vine et al., 2013, 2017). Visual guidance by external focus of attention, that is, focusing on the target leads to a more effective golf swing than technical training of the postures and limb movements composing a golf swing. Indeed, in contrast with technical training, QE training of golf putting leads to improved putting accuracy as well as to a reduction in heart rate and muscle activity at the moment of impact (Moore et al., 2012). Thus, it is believed that instructing athletes by directing their perceptual linkages outward to the visual field supports higher task performance as well as leads to more expert-like movement and physiology.

QE research opens a fascinating vantage point on the perceptual linkages anchoring organisms to their visual contexts and the synergies transforming visual information into action (e.g., Kelty-Stephen et al., 2020; Mangalam, Lee, et al., 2021; Profeta & Turvey, 2018). Research on QE has mostly examined the visual-cognitive bases for the relatively more extraverted gaze into the task context (Panchuk & Vickers, 2006; Rienhoff et al., 2016; Vickers, 2011), and the relationship of QE to the body, the brain, and the nervous system has received relatively less attention (Davids & Araújo, 2016; Renshaw et al., 2019; Shine et al., 2018). To fill this gap, a ‘postural-kinematic hypothesis’ has begun to complement the canonical ‘visual hypothesis’ that QE was a primarily visual-cognitive mechanism (Gallicchio & Ring, 2020). So far, this postural-kinematic hypothesis emphasizes only slower movement times, that is, longer duration offering the foundation for longer fixations. This postural-kinematic hypothesis is a constructive first step. In this study, we aim to embed more of the postural dynamics into this hypothesis, beyond simply movement duration. Additionally, instead of reporting on gaze measurements, we will use QE as a training manipulation and focus on investigating how postural sway supports the effect of the training.

### 1.2. Optic flow and postural sway can support the quiet eye strategy

A crucial consideration for postural-kinematic contributions to visually guided tasks is optic flow (Stoffregen, 1985, 1986; Stoffregen & Riccio, 1990). Amidst the ‘noisy’ fluctuations of the athlete’s body, quieting the eyes might leave fewer saccadic interruptions to the coupling between optic flow and postural sway. Optic flow consists of the differential expansion, contraction, or rotation of visible objects and surfaces as an individual moves about. It is specific to both the individual’s movement and the contextual layout of objects and surfaces. Indeed, virtual visual displays that impose artificial optic flow also engender anticipatory behaviors (e.g., steering, braking) as if the objects and surfaces entailed by the optic flow were out there—not just for humans but for other animal species and even robots (Serres & Ruffier, 2017). Although most prominent in locomotion-related contexts, postural sway is no less capable of generating informative optic flow in non-locomotion contexts (Stoffregen et al., 2005). Because research on QE has mostly focused on the fixation of the eye gaze, QE’s relationship to optic flow remains a critical knowledge gap (de Oliveira, 2016).

On the face of it, the preceding arguments might ring hollow at first. Certainly, optic flow requires movement, and it could seem wholly unrelated to the issue of visual fixations supporting wholly internal processing of incoming information into a motor program for the body’s subsequent control. This gap between QE and optic flow follows closely from ongoing controversy in the literature between ecological vs. information-processing explanations. Optic flow has ecological roots in Gibson’s (1979) disavowal of information-processing. Those roots could potentially raise fears that this work might disregard the extensive, diligent work on the information-processing account of QE. Indeed, a rare mention of ‘optic flow’ in the information-processing approach to QE weds it firmly to ‘the teachings of Gibson,’ so concerned with invariance as to ‘treat every gaze as the same,’ and reluctant to respect the fine-grained dynamics of gaze (Vickers, 2009, p. 286). We make full disclosure: the present use of multifractal modeling of postural access to optic flow is inspired by Gibson’s appeal to flows and nestings these flows across many scales (Kelty-Stephen & Wallot, 2017). However, we come to this dialogue with first-hand experience examining the rich heterogeneity of gaze dynamics (e.g., Kelty-Stephen & Mirman, 2013; Stephen & Mirman, 2010) and with such interest in cognitive process as for Gibsonians to openly doubt that some of us could be Gibsonians (Raja & Anderson, 2019). Indeed, our interest in how exploratory behaviors might change perceptual response sets us firmly outside one of the stricter readings of Gibsonian invariance (e.g., Turvey, 1977).

Rather than digging into old trenches, we think that optic flow has a theoretical reach that might pick up where this information-processing account of QE remains incomplete. The theoretical reach of optic flow has grown well beyond Gibson’s original intent and usage. For instance, we have already cited cases of optic flow that would have been startling through the strictly Gibsonian lens: information-processing robots use optic flow as well as Gibson’s organisms might (Serres & Ruffier, 2017), and optic flow does not require locomotion but can thrive on postural sway (Stoffregen et al., 2005). In the spirit of making the so far dialogue between ecological and information-processing approaches more constructive, the preceding two examples suggest that optic flow might be much more subtle (e.g., as in standing ‘still’) but also perfectly amenable to information processing (e.g., as in supporting computational models for controlling action).

Meanwhile, without optic flow to support it, QE’s information-processing accounts are incomplete and unmoored from their own explicit requirements and proposals. According to a recent summary, the information-processing view of QE is that the eye’s stillness provides time and attentional resources useful for the preprogramming of movements used in the aimed behavior (Vickers, 2009). This point is defensible but incomplete: visual-cognitive processes benefit only partially from a motor program’s stipulation. Even non-ecological explanations have become newly interested in emergent control parameters that sit outside of any simple stipulation by an internal motor program (Vecera et al., 2014). Visual attention is most effective when it is most task- and context-sensitive. The generic usefulness of QE points to some long-unanswered questions. Are the motor programs implicated in QE flexible and vast enough to accommodate both bodily degrees of freedom and contextual ones? Or are these motor programs elegantly minimal enough to leave room for non-programmed factors to support? Neither option is unappealing on its face, but they both reflect an underdeveloped delivery on the information-processing account’s proposals about QE. Yes, longer looking times can support stronger, more enriched motor programs (e.g., Witt & Proffitt, 2008). However, it is not always so simple: either looking time is not always a reliable predictor of motor planning, or the motor program is flexible enough to be rewritten on the fly during the program’s execution (Hajnal et al., 2014, 2016).

Specifically, just as it neglects the finer print about its motor programs, QE’s information-processing account has also neglected its claims about the role of action. The information-processing account offers QE a bidirectional link between gaze, attention, and action (Vickers, 2009). However, whereas this view may respect bidirectionality between attention and gaze (i.e., attentive search for information and looking to what cues attention), the action appears strictly as the outcome passively inheriting whatever the preprogramming entails and only following motor commands. If motor programs generate the consequent swings, then the information-processing approach has yet to elaborate upon two points: first, what bodywide movement coordination supports the postural control allowing the steady visual fixation and, second, how the preprogramming contributes to the externally focused movement. For instance, especially as participants repeat the aiming tasks in front of potentially the same experimental display, the evidence suggests more variety in expert movements than sameness, mainly as they engage visually in the external focus of attention (Morrison et al., 2016). This point raises some questions. If QE is merely the same sort of fixation every time, irrespective of movement, then why should programs carrying similar, static information give rise to varied bodily execution of the program? If the motor programs that QE supports vary, what is the possible source of variability? In the former case, the programs might underspecify the movements, and in the latter case, the programs would seem to embody more variety than the fixity of the visual scene. The whole issue entailed by the Bernsteinian ‘degrees of freedom’ concern is that there is no clear programmatic path from intention to bodily execution. Suppose action did participate in the same bidirectional relationship that information-processing approaches to QE observe between attention and gaze. In that case, there might be a way forward, allowing for action to participate in the tuning of programs to ongoing bodily postures and physiological changes throughout repeated behaviors.

To summarize, the gap between QE and optic flow is critical because the information-processing approach aligns the fixity of the eyes with the accuracy of aimed behavior. However, there is a little account of the body between the fixations and the accurate behaviors. This omission of the body is immensely curious and potentially problematic because movement exhibits rich variability. It has been difficult for all proposed motor programs to specify accurate goal-directed behavior (Feldman, 2018) and no less difficult when eye movements are implicated (Feldman & Zhang, 2020). Moreover, if we can leave off old associations and let optic flow continue to take on shapes beyond the original Gibsonian imaginings, we might reinstate the body to an account of QE without any threat of thwarting or disregarding prior art. We aim to rekindle this dialogue between cognitive psychology and ecological psychology, but we aim to do so without requiring disavowal of any mutually opposed pre-theoretical commitments.

The present work aims to replicate the beneficial effects of QE on putting performance, and more importantly, to understand how eye height and optic flow might support QE. We used a single session in the style of Moore et al.’s (2012) golf-putting training paradigm to test for rapid attunement to changes in information from optic flow and postural sway. Optic flow centers on an external focus of control, aligned with their eye height (e.g., Warren, 1984), and when we perturb eye height, we destabilize optic flow, and the organism needs to learn new relationships between bodily movement and movement of visible surfaces (Frenz et al., 2003). So, we aimed to test whether perturbing the optic-flow parameter of eye height perturbs the effect of QE training in golf putting more so than technical training. In non-golf contexts, for instance, manipulating eye height by adding wooden blocks to participants’ shoes alters participants’ affordance judgments significantly, leading to an initial underestimation of the optimal climbing height for a stair and sitting height for a chair (Warren, 1984). However, participants adjust to the new eye height and make more accurate decisions shortly afterward (Mark, 1987; Mark et al., 1990). Hence, we expected that adding wooden blocks—(henceforth, ‘clogs’, to distinguish from ‘blocks’ of experimental trials)—to the feet might upset putting accuracy in the QE-trained participants initially, but that this accuracy would rebound as participants adjust to new eye height (Rienhoff et al., 2016).

### 1.3. Multiscaled organization of postural sway entails a multifractal structure

Besides manipulating eye height, our expectations about the optic-flow foundation for QE also included a hypothesis about postural sway moderating effects of perturbed eye height. If optic flow could support QE, we expected that QE-trained response to eye height perturbation would depend on postural sway. Indeed, in the affordance research on reattunement to eye-height perturbations, postural sway supported the adaptation of the eye-height relationship to judgments of optimal sitting height (Mark et al., 1990). However, the exact aspects of sway supporting adaptation have been unclear. It was not simply that more sway facilitated this adaptation because, for instance, an ‘awkward stance’ requiring more compensatory postural adjustments did not support visual adaptation as much as comfortable standing did. An additional challenge in identifying adaptation-supporting aspects of sway could be that sway unfolds across multiple timescales. At longer scales, QE training itself might influence sway by training an explicit use of visual information. At shorter timescales, adding clogs might destabilize posture initially, and then at the shortest timescales, sway might be implicated in adapting to this perturbation of eye height. Hence, judicious accounting of sway’s contribution to visual adaptation requires an appreciation of sway’s multi-scaled organization.

Then again, the multi-scaled organization of sway may itself reveal a key to predicting—if not explaining—how the movement system adapts its visually-guided actions. Interactions across multiple timescales entail a ‘multifractal’ structure in sway. Multifractality refers to multiple (i.e., ‘multi-’) relationships among fluctuations and timescales, with these relationships following power laws with potentially fractional (i.e., ‘-fractal’) exponents. Interactions across timescales engender a nonlinear patterning of fluctuations that multifractal analysis can diagnose and quantify (Ihlen & Vereijken, 2010). Specifically, multifractal analysis estimates a spectrum of fractal exponents whose width *W*_MF_ quantifies “multifractality.” When we compare measured series’ *W*_MF_ to spectrum widths of a set of best-fitting linear models (‘linear surrogates’) of those series, multifractal evidence of nonlinearity manifests as a significant difference between *W*_MF_ for the original series and the average *W*_MF_ for linear surrogates. The *t*-statistic, *t*_MF_, comparing the original series’ *W*_MF_ to the surrogates’ *W*_MF_ serves as an estimate of what we call ‘multifractal nonlinearity’ (i.e., how much multifractality in the original series is due to nonlinear interactions across timescales).

### 1.4. Multifractal nonlinearity predicts perceptual responses to perturbations of task constraints

Multifractality, *W*_MF_, in postural sway is a robust predictor of perceptual responses in both visual and haptic media (Doyon et al., 2019; Hajnal et al., 2018; Teng et al., 2016). Multifractal nonlinearity, *t*_MF_, in postural sway predicts perceptual responses as well, not just in the case of nonvisual judgments of length and heaviness of manually-wielded objects (Mangalam & Kelty-Stephen, 2020) but also in visuomotor tasks (Bell et al., 2019; Booth et al., 2018) and in response to perturbations, such as those due to prismatic goggles in a visual aiming task (Carver et al., 2017). Now, we aimed to test whether *t*_MF_ could predict the responses to perturbations of eye height in QE-trained golf putting.

We expected that postural sway most adaptive to the perturbation of eye height would show relatively narrow multifractal spectra that would still differ significantly from those of the surrogates. So far, postural sway shows relatively narrower multifractal spectra (i.e., smaller *W*_MF_) for younger participants (Munafo et al., 2016), those less likely to experience motion sickness (Koslucher et al., 2016), and those focusing on more proximal surfaces in optic flow (Li et al., 2020). Hence, greater *W*_MF_ in postural sway might counteract effects of QE. All these findings are qualified by a failure of *W*_MF_ to correlate with canonical measures of sway like standard deviation. However, postural stability is only ambiguously related to the standard deviation of sway, requiring neither too much nor too little variability in sway (Chen et al., 2008; Rajachandrakumar et al., 2018). Meanwhile, both empirical and theoretical work suggests that *t*_MF_ is more closely related to postural stabilizing. Postural stabilization following a perturbation to the base of support shows a rapid reduction of *t*_MF_ (Kelty-Stephen, 2018). Theoretical simulation also predicts that random perturbations to individual scales of a nested system lead to narrower multifractal spectra that still differ from linear surrogates’ spectra (i.e., smaller *W*_MF_ and smaller-yet-significant *t*_MF_; Lee & Kelty-Stephen, 2017). Hence, adapting to the posturally perturbed visual information might involve exhibiting smaller-yet-significant *t*_MF_.

### 1.5. Golf-putting task and hypothesis

The participants were instructed to take golf putts towards a circular, hole-sized target painted on a green putting surface in the laboratory—over four blocks of ten trials each. One group received QE training, and the other group received technical training specifying how to move their limbs to make the golf putt. Wooden clogs were attached to the feet of all participants to perturb their eye height. Putting performance was treated continuously in terms of unsigned radial distance from the target (i.e., unsigned radial error).

#### 1.5.1. Hypothesis-1: Multifractal fluctuations in postural sway will lead an eye height perturbation to interrupt QE-trained performance more so than technical-trained performance

We hypothesized that the eye-height perturbation due to clogs would lead to lower putting accuracy characterized by higher unsigned radial error following QE training than technical training. We predicted that the clog’s effects on QE would depend on the multifractal nonlinearity, *t*_MF_, of sway above and beyond any effects of multifractality (i.e., *W*_MF_) alone (Hypothesis-1a). We also predicted that lower *t*_MF_ of sway would be associated with more adaptive responses to the eye-height perturbation, leading to gradual increase and subsequent decay of radial error across trials within the blocks with the clogs on (Hypothesis-1b).

## 2. Methods

### 2.1. Participants

Twenty undergraduate students (11 women, *mean*±1*s.d.* age = 19.5±1.2 years, all right-handed, Oldfield, 1971) with normal or corrected vision participated in this study. All participants provided written informed consent approved by the Institutional Review Board (IRB) at Grinnell College (Grinnell, IA).

### 2.2. Procedure

In an hour-long session, the participants completed a single putt on each of 40 trials. Prior to the training trials, half of the participants received QE instructions (Table 1, left column) while the other half received technical putting instruction (Table 1, right column), replicating Moore et al.’s (2012) methodology. The technical instructions mirrored the QE instructions but did not describe the eye-related behaviors, hence mimicking standard golf instruction. After verbally confirming that they understood the instructions, the participants watched a video demonstrating a QE-specific putting stroke (https://www.youtube.com/watch?v=lfWuL1klbsY&feature=youtu.be) or a standard putting stroke (https://www.youtube.com/watch?v=HOYZn4MYmDI). The researcher directed the QE-strained participants to the key features of the golfer’s gaze control and the technically-trained participants to the putting mechanics.

**Table 1.**
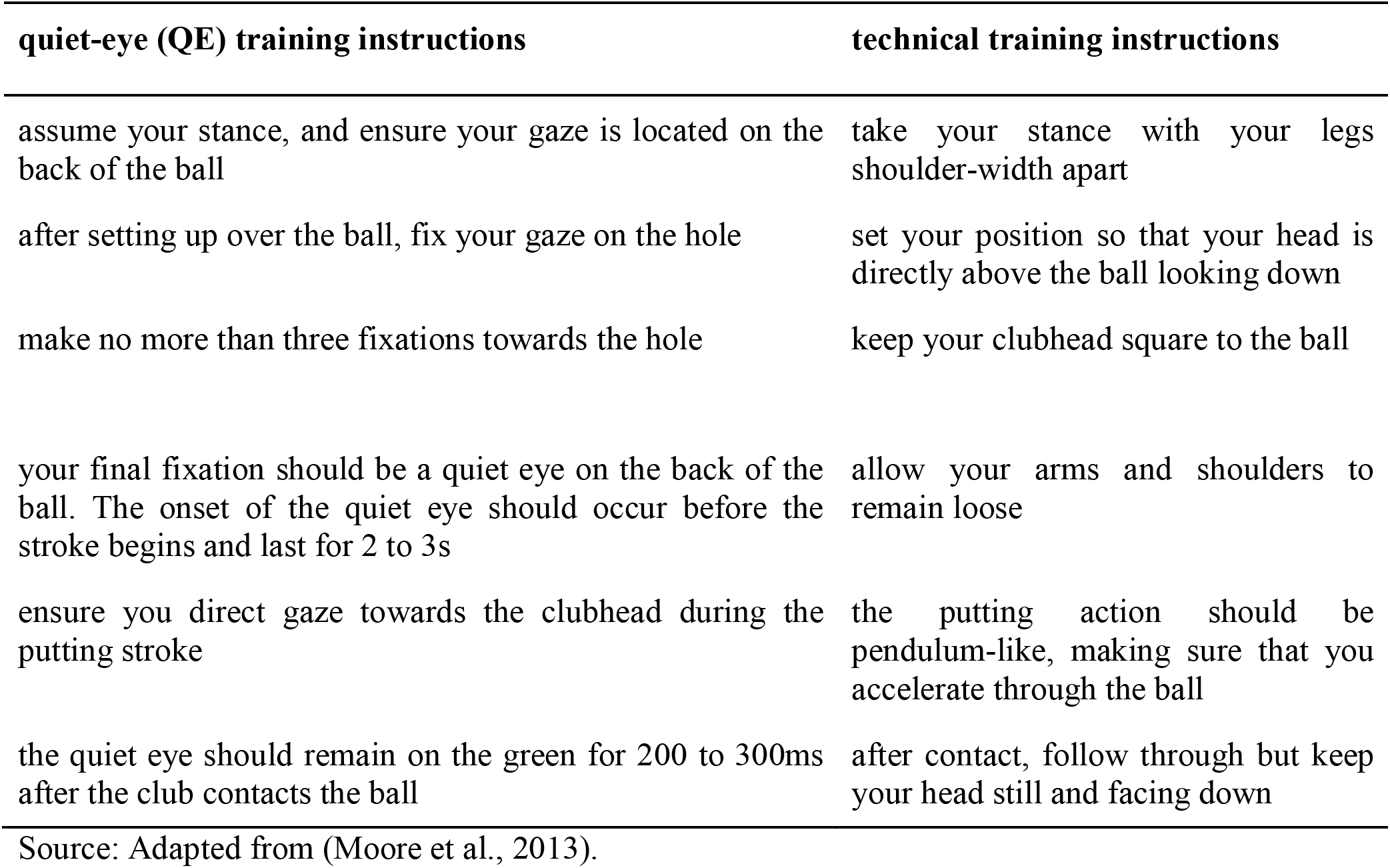
Training instructions given to the quiet-eye and technical groups during the training phase.

The participant performed the golf putts on a 6×15 feet standard indoor All Turf Matts™ artificial turf mat using a 1.68-inch diameter PING Sigma 2 Anser Stealth™ putter and regular-sized white golf balls. The participants completed straight putts ten feet away towards a 2.5-inch diameter circular orange dot, designed to fit the golf ball. This format allowed for more stringent measurement and gave the participants a more focused target.

The participants completed 20 training trials (treated as two blocks of 10 trials per block in regression modeling). Following a short 5-minute break, the participants completed two blocks of ten test trials per block (i.e., 20 test trials total) in the absence of instructions. The participants were randomly assigned to complete either their first or second block of ten trials while wearing 5-cm-raised wooden blocks (i.e., clogs) attached to their shoes (they completed the other block of ten test trials under the same conditions as the 20 training trials). Before the testing began, the participants were allowed to walk around and adjust to the clogs to reduce fall risk.

### 2.3. Measures

The task performance was defined in terms of the radial error (the distance from the target at which the ball stopped, in feet) and the number of putts successfully ‘holed.’ Because the target was not a real hole, the putt was counted as holed in either of two events: the ball came to rest within the orange dot on the floor or within 2 feet directly beyond the hole (Cooke et al., 2010; Vine et al., 2011). Postural sway was measured using a smartphone with the Physics Toolbox Suite application (http://vieryasoftware.net) strapped to the participants’ lower back. The app recorded the acceleration of the torso along *x*-, *y*-, and *z*-dimensions at 100 Hz. An experimenter manually started and stopped recording for each putt. Recording began after the participant set up over the ball and stopped once the ball stopped moving.

### 2.1. Multifractal analysis

#### 2.4.1. Multifractal spectrum width of accelerometer displacement series

For each putt, we computed an accelerometer displacement series for multifractal analysis as follows. The accelerometer data encodes an *N*-length series of accelerations of the torso along *x*-, *y*-, and *z*-dimensions centered on an origin on the smartphone. A displacement series of length *N* − 1 was computed by: 1) taking the first differences for the variable (e.g., if we use the letters *i* and *j* to indicate individual values in sequence, *x_diff_* (*j*) = *x*(*i*) − *x*(*i* − 1) for each uth value of *x*, for 2< *i* < *N*, 1< *j* = *i* − 1 < *N* − 1, 2) squaring and summing the differences of each dimension’s displacements (i.e., 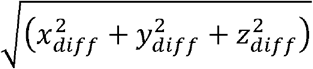), and 3) taking the square root of this sum (i.e., 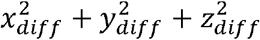).

Multifractal spectra *f*(*α*) were estimated for each accelerometer displacement series, using Chhabra and Jensen’s (Chhabra & Jensen, 1989) method. The first step was to partition the series into subsets (or ‘bins’) of various sizes, from four points to a fourth of series’ length (Fig. 1, left). The second step was to examine how the proportion of displacements within each bin changed with bin size or timescale (Fig. 1, right). The third step was to estimate two distinct exponents: an exponent *α* describing how the average proportion grows with bin size and an exponent *f* describing how Shannon’s (Shannon, 1948) entropy of bin proportion changes with bin size. The later steps were all iterations of this third step, calculating a mass *μ* by applying an exponent *q* to bin proportions and using *μ*(*q*) to weight proportions selectively according to proportion size. *q* greater than 1 weights larger-proportion bins, and *q* lesser than 1 weights smaller-proportion bins (Fig. 2, left). *q*-based mass *μ*(*q*)-weighting generalizes single *α* and *f* estimates from earlier steps into continual *α*(*q*) and *f*(*q*)— with *α*(*q*) describing how mass-weighted average bin proportion changes with bin size and *f*(*q*) describing how Shannon’s entropy of bin masses *μ*(*q*) with bin size (Fig. 2, top right).

**Figure 1.**
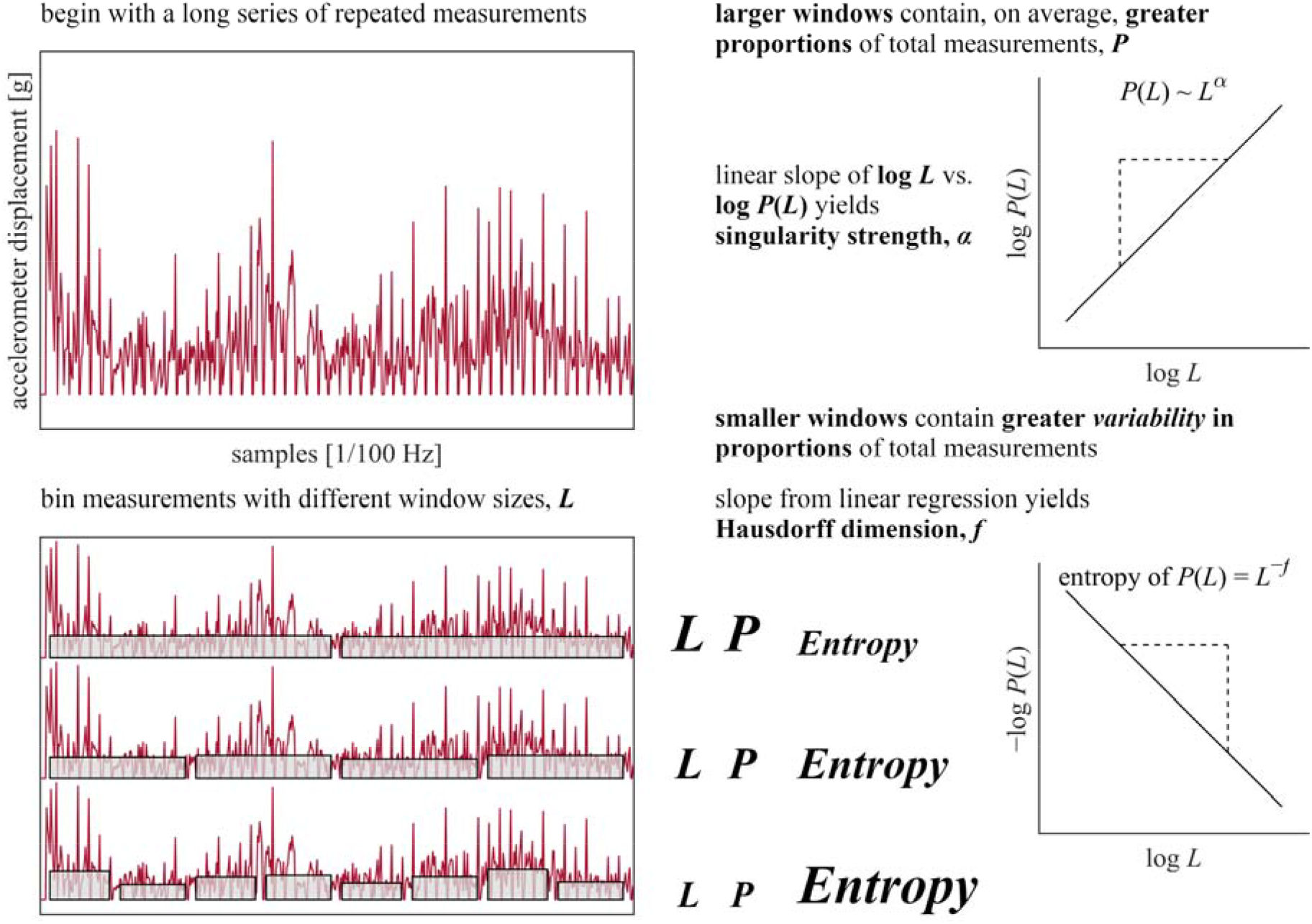
Schematic of Chhabra and Jensen’s (Chhabra & Jensen, 1989) method used to estimate multifractal spectrum *f*(*α*) (Part-1). See Section 2.4.1. for further details.

**Figure 2.**
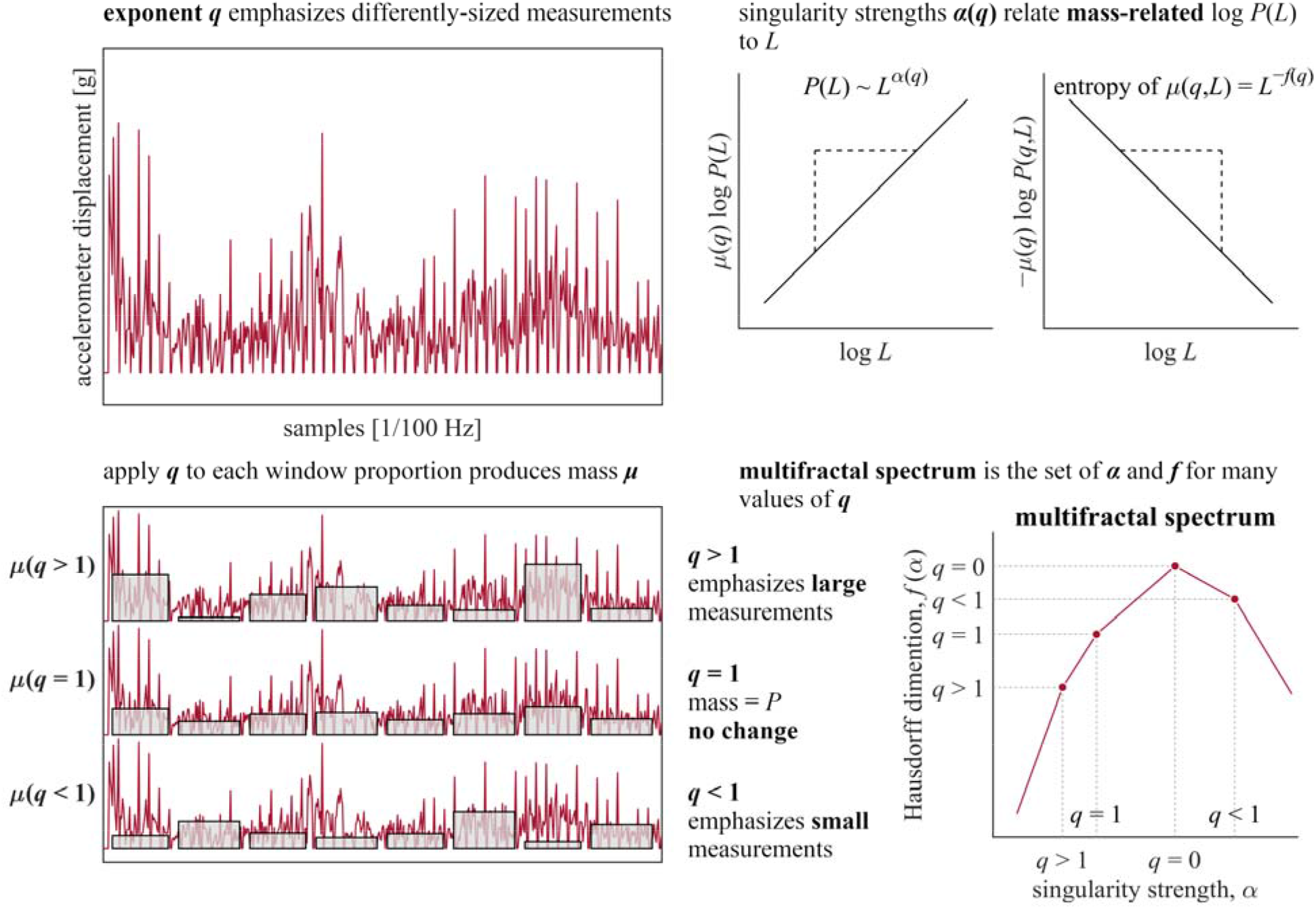
Schematic of Chhabra and Jensen’s (Chhabra & Jensen, 1989) method used to estimate multifractal spectrum *f*(*α*) (Part-2). See Section 2.4.1. for further details.

Multifractal analysis quantifies heterogeneity as variety in *α*(*q*) and *f*(*q*). The ordered pairs (*α*(*q*), *f*(*q*)) constitutes the multifractal spectrum (Fig. 2, bottom right). We included *α*(*q*) and *f*(*q*) only when mass-weighted proportions were defined and when both of their corresponding relationships to bin size correlated at *r* > 0.995 on logarithmic plots, for bin sizes 4,8,12,…*N*/4 and −300 < *q* < 300, where *N* is the number of samples in the series.

#### 2.4.2. Surrogate comparison

Multifractal spectrum width is sensitive to nonlinear interactions across timescales as well as to linear temporal structure. Hence, evidence of nonlinear interactions across timescales requires comparing the original series’ multifractal spectrum width *W*_MF_ to those computed for a sample of linear surrogate series (Fig. 3). These linear surrogate series retain the same value as the original series but in a different order that preserves the original series’ linear autocorrelation but destroys original sequence (i.e., iterative amplitude-adjusted Fourier-transform or IAAFT (Ihlen & Vereijken, 2010; Schreiber & Schmitz, 1996); Figs. 4*a* to *d*). Comparing *W*_MF_ of the original and surrogate series (Figs. 4*e*, *f*) produces a *t*-statistic *t*_MF,_ a measure of nonlinear interactions across timescales that is standardized across series with differing linear temporal structure. *t*_MF_ was computed for each original series.

**Figure 3.**
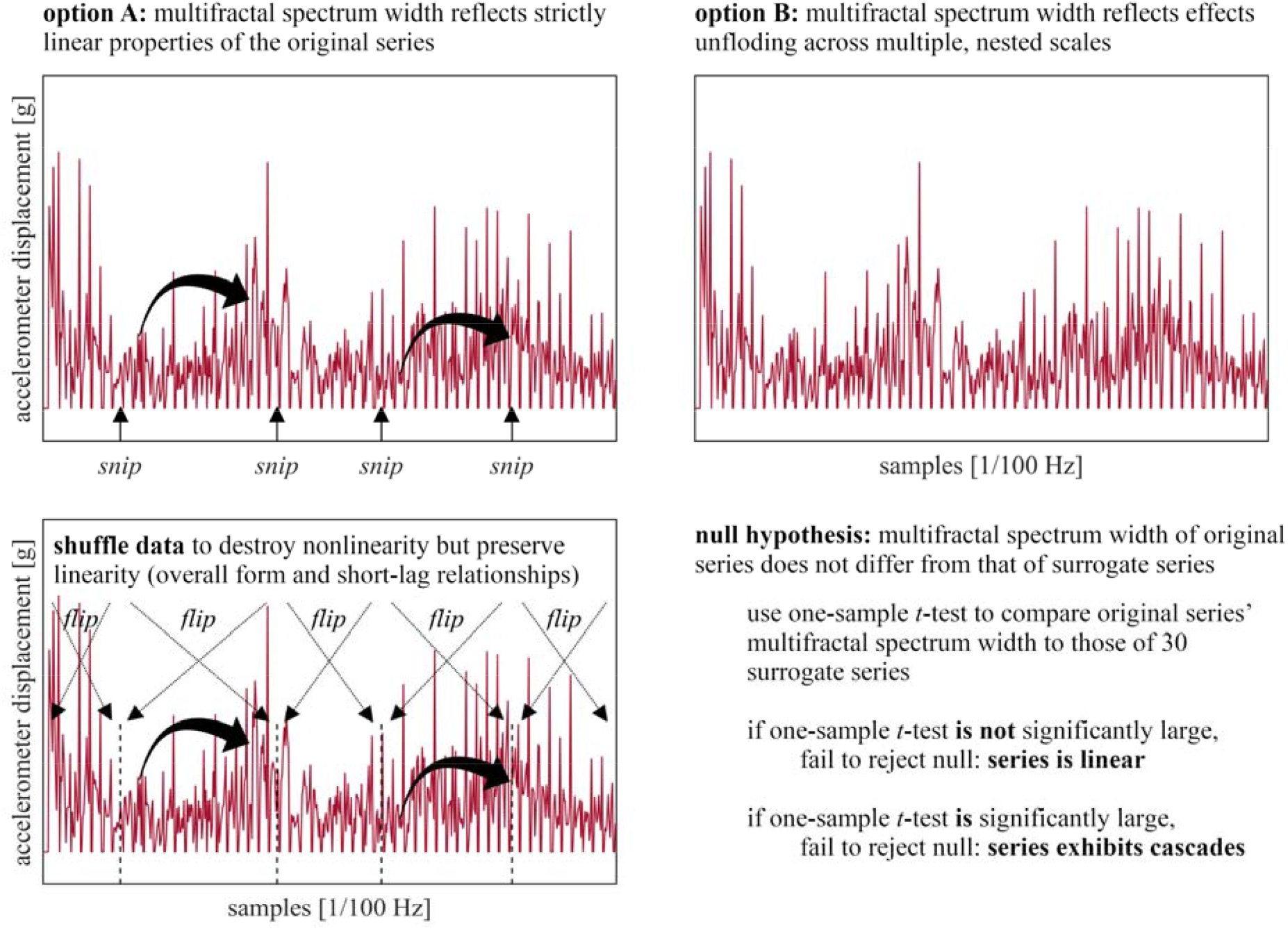
Surrogate analysis used to identify whether a series exhibits multifractality due to nonlinearity (cascades). See Section 2.4.2. for further details.

**Figure 4.**
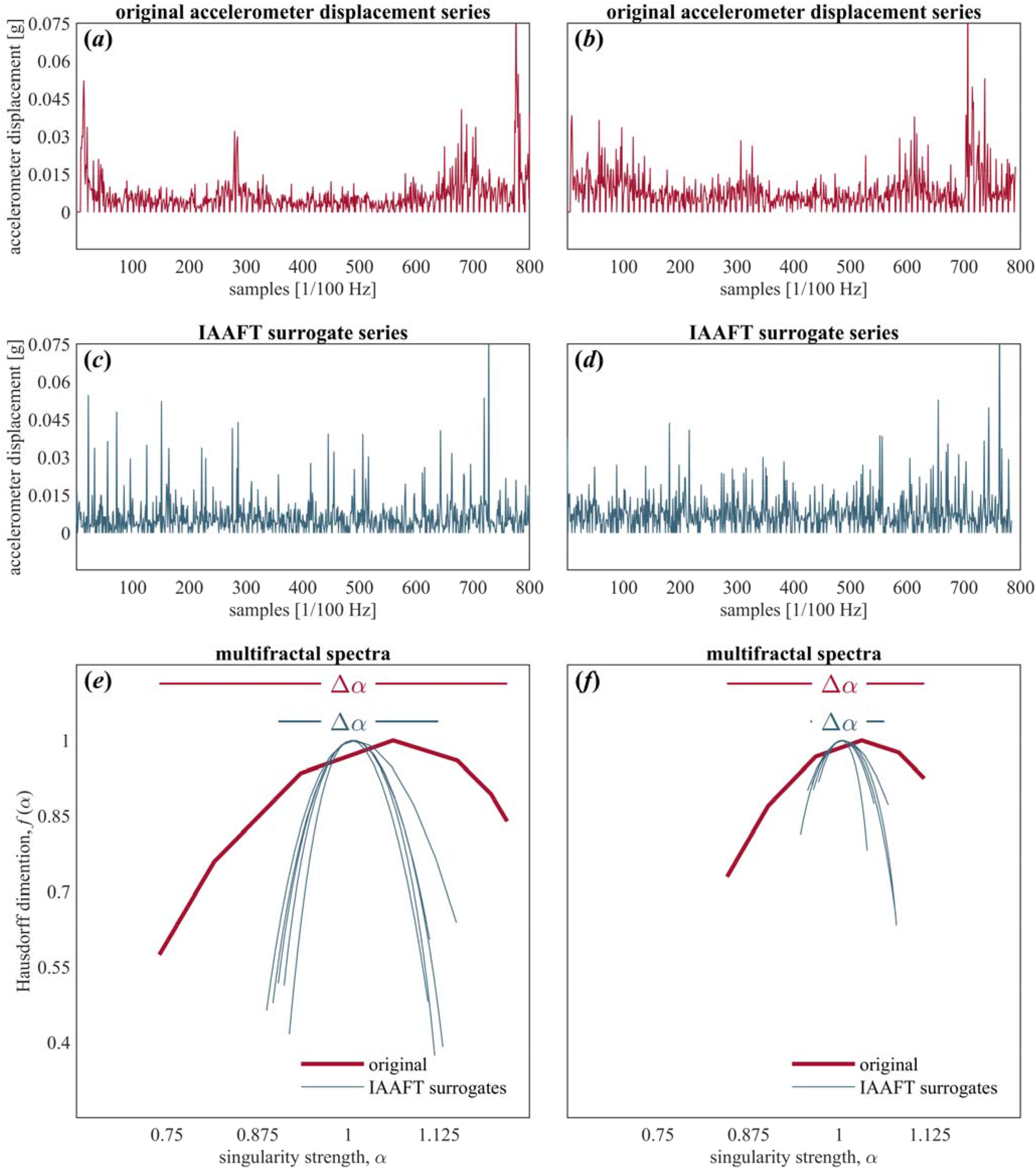
Multifractal analysis. (*a*, *b*) Representative original accelerometer displacement series. (*c*, *d*) Representative IAAFT surrogate series of the original accelerometer displacement series in *a* and *b*. (*e*, *f*) Multifractal spectrum (*f*(*α*), *α*) of original series in *a* and *b*, and those of their five representative IAAFT surrogates.

### 2.5. Mixed-effects modeling

The data were submitted to two kinds of mixed-effect modeling, namely, logistic and Poisson, using the glmer() function in *R*-package *lme4* (Bates et al., 2007). The subsequent subsections provide details of the predictors and the models.

#### 2.5.1. Effects of time

The effects of time were modeled using trial number within a block (trial_block_), block number (block), and interactions of trial number with block number (trial_block_ × block), and trial number across blocks (trial_exp_). The model used orthogonal polynomials of each of both predictors up to 3^rd^ order (i.e., cubic). The primary reason for the polynomial terms was to reveal how and when the perturbation of wearing clogs might manifest and consequently subside. Cubic polynomials allow the simplest way to accomplish this goal by allowing a peak and subsequent decay to a flat, quiescent portion as part of the same continuous function. Additionally, unlike the quadratic function, the cubic peak is not required to be symmetric and centered on the midpoint. Hence, the cubic function controls for any nonlinearities in the normal course of trials without such perturbations. Whereas block is a linear effect across blocks, including trial_exp_ in the model allowed controlling for nonlinear change across blocks.

There are two important features to note about the treatment of time in this model. First, the Poisson model encoded both classifications of trial numbers (i.e., trial_exp_ and trial_block_) in terms of an orthogonal 3^rd^-order polynomial, using the function ‘poly’ in *R*, to include the linear, quadratic and cubic growth of trial number with all three terms phase-shifted so as to be uncorrelated with one another. Second, the lack of model convergence limited the appearance of block, such that block was neither treated nonlinearly nor in interaction with perturbation. Attempting to include the orthogonal 3^rd^-order polynomials for block number led to a failure of model convergence, so block only appeared as a linear term. The model also failed to converge when it included interactions of block and perturbation because these two terms were strongly collinear.

#### 2.5.2. Effects of manipulations

Manipulations included quiet-eye instruction, QE (QE = 1 for receiving QE instruction, and QE = 0 for not receiving QE instruction) and perturbation of eye height, perturbation (perturbation = 1 for all ten trials during the block when participants wore the clogs, and perturbation = 0 for all other trials).

#### 2.5.3. Sway: Linear predictors

The accelerometer data encodes accelerations of the torso along *x*-, *y*-, and *z*-dimensions. Any single posture corresponds to a specific point in this 3D space. Hence, sway corresponds to the deviation of this 3D acceleration around the mean posture. The present analyses encoded this variability as *MSD*_acc_ or *RMS*_acc,_ depending on which predictor supported a convergent model, along with mean and standard deviation of 3D displacement (*Mean*_disp_ and *SD*_disp_, respectively).

#### 2.5.4. Regression modeling

A mixed-effects Poisson regression modeled the radial error (in inches) of all misses, with makes as defined above treated as having zero radial error. Predictors included QE × perturbation × trial_block_ × *t*_MF_ to our hypothesis by specifically addressing trials within each block, QE × trial_block_ and QE × trial_exp_ to control for differences in learning due to QE across a linear progression of block; a nonlinear function of trials, QE × perturbation × *W*_MF_ × *t*_MF_, to control for the effects of multifractal spectral width and its relationship with *t*_MF_; and *MSD*_acc_ × *Mean*_disp_ × *SD*_disp_ to control for the effects of linear description of sway. Modeling included lower-order interactions and component main effects (e.g., Allison, 1977).

## 3. Results

### 3.1.1. Non-multifractal effects on radial error

This section details the non-multifractal effects described in Supplementary Table S1 and effects preliminary to the test of our hypothesis in Table 2. This model of radial error found a significant main effect of QE training, specifically, a negative effect (*b* = –4.40×100, *s.e.m.* = 4.81×10^−1^, *p* < 0.0001) indicating that QE did promote greater accuracy in terms of proximity of the ball’s final position to the hole. Radial error grew linearly and quadratically across trials but showed negative cubic variation with trial—both for trials across the experiment and within a block (Fig. 5a). The QE strategy prompted steady decay of error across the experiment, canceling out the linear and cubic pattern of error over trials within a block (Fig. 5b). The perturbation of eye height elicited no main effect but increased the growth of error over trials within a block, with the QE instruction accentuating this nonlinear growth of error later in the block. The higher-order interaction *MSD*_acc_ × *Mean*_disp_ × *SD*_disp_ and its main component effects and lower-order interactions showed mostly negative effects (only two of the two-way interactions showed positive coefficients).

**Table 2.**
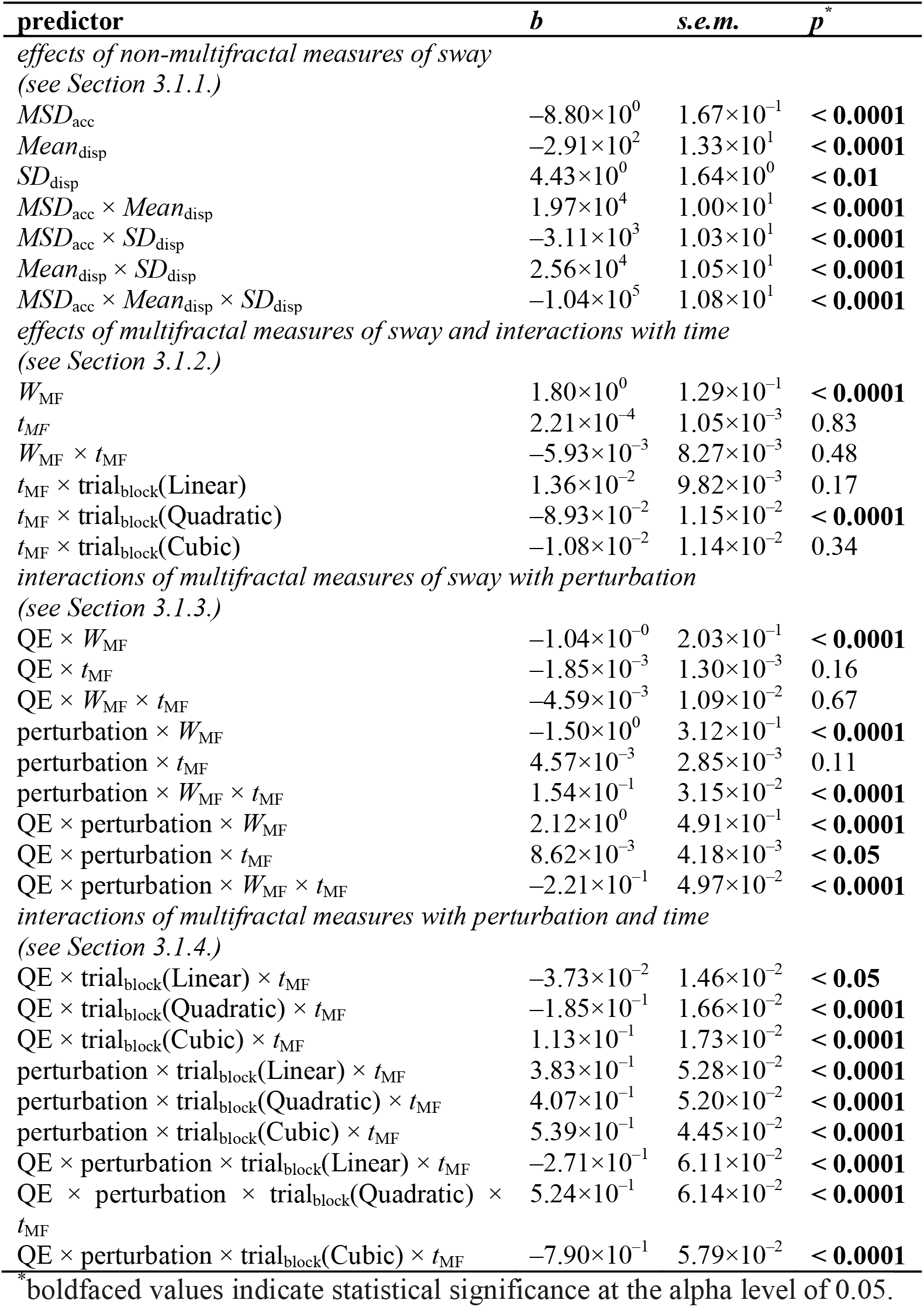
Subset of Poisson regression model including main and interaction effects non-multifractal and multifractal measures of sway, perturbation of eye height, and time

**Figure 5.**
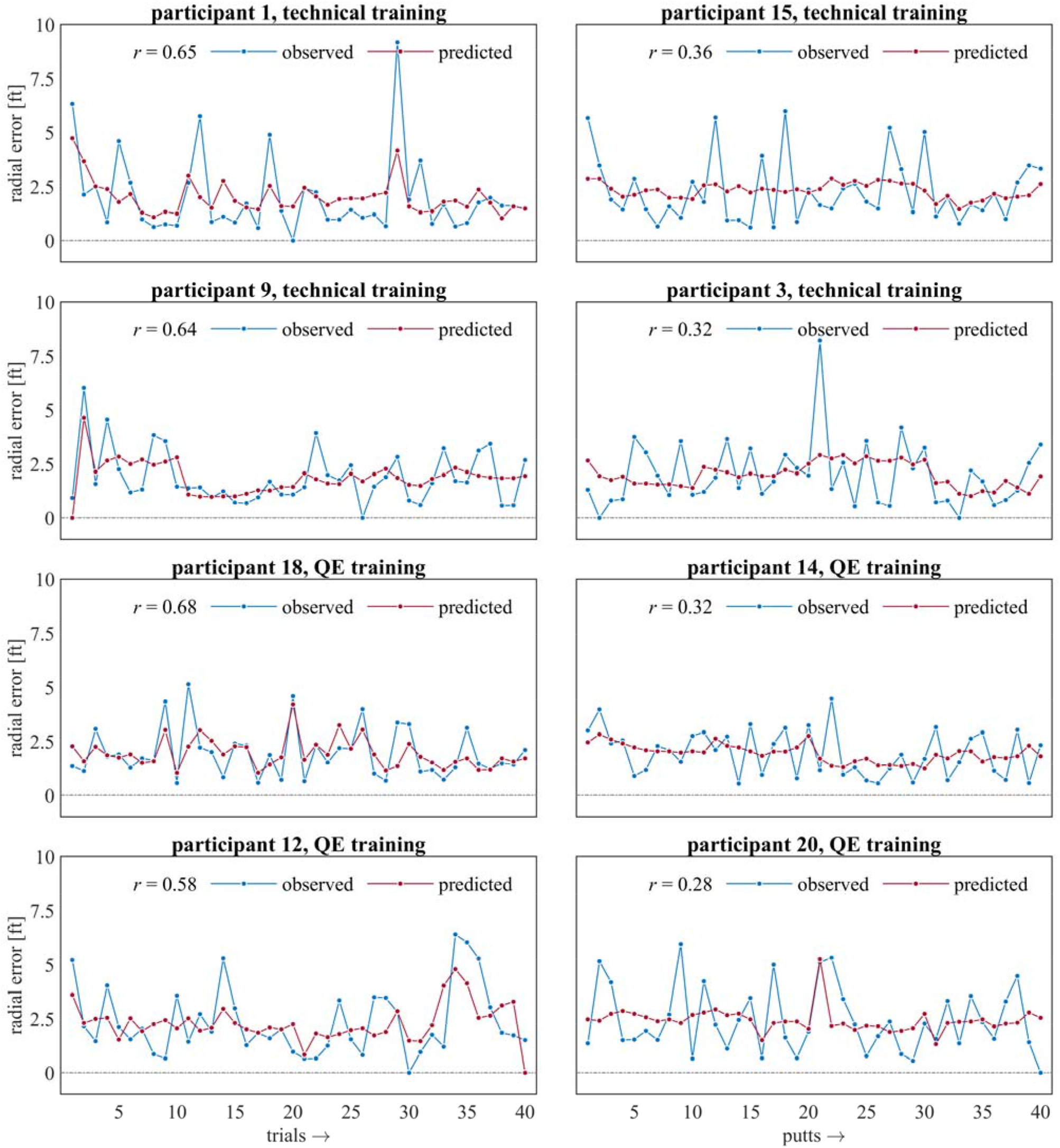
Trial-by-trial model predictions of radial error for the two participant groups. (*a*) Technical-trained group. (*b*) QE-trained group.

### 3.1.2. Total multifractality of sway, *W*_MF_, increased radial error, but multifractality due to nonlinearity, *t*_MF_, attenuated radial error at the beginning and the end of blocks

No matter what portion of multifractality is attributable to linear or nonlinear structure, the total width *W*_MF_ of original series’ multifractal spectra is associated with greater radial error (*b* = 1.80×10^0^, *s.e.m.* = 1.29×10^−1^, *p* < 0.0001). We did not find any main effect of multifractality due to nonlinearity, *t*_MF_, nor any interaction of *W*_MF_ × *t*_MF_. However, the negative effect for *t*_MF_ × trial_block_ (Quadratic; *b* = – 8.93×10^−2^, *s.e.m.* = 1.15×10^−2^, *p* < 0.0001) indicated an association between multifractality due to nonlinearity with a significant reduction of the positive quadratic profile for radial error over trials within a block (i.e., trial_block_(Quadratic); Section 3.1.1.).

### 3.1.3. Testing Hypothesis-1a: Perturbation of eye height increased radial error only in combination with QE training, with greater multifractality due to nonlinearity *t*_MF_, or with both

The multifractality of sway promoted the tendency of QE to increase perturbation-related radial error. Perturbation showed a significant effect through the interaction of perturbation × *W*_MF_ (*b* = –1.50×10^0^, *s.e.m.* = 3.12×10^−1^, *p* < 0.0001) canceled out the positive effect of *W*_MF_ on error (Section 3.1.2), suggesting that the perturbation reduced the error-eliciting effect of *W*_MF_. The interaction QE × *W*_MF_ reduced error due to multifractality by roughly half (*b* = –1.04×10^0^, *s.e.m.* = 2.03×10^−1^, *p* < 0.0001), suggesting that QE promoted greater accuracy but only in the absence of perturbation. However, the higher-order interaction with multifractality indicates that QE increased the error attributable to the perturbation. For instance, the QE × perturbation × *W*_MF_ (*b* = 2.12×10^0^, *s.e.m.* = 4.91×10^−1^, *p* < 0.0001) reverses the apparently beneficial error-reducing interaction effect of perturbation × *W*_MF_. Hence, QE reinstated the error due to total multifractality (Section 3.1.2), and it did so only in the face of the perturbation.

Above and beyond the effects of *W*_MF_ in moderating the interaction effect of QE × perturbation, *t*_MF_ promoted the error-increasing effect of perturbation. Specifically, besides the moderating effect of QE on the effect of perturbation × *W*_MF_, perturbation elicited even greater radial error with increases in *t*_MF_ (perturbation × *W*_MF_ × *t*_MF_: *b* = 1.54×10^−1^, *s.e.m.* = 3.15×10^−2^, *p* < 0.0001). QE did reverse this effect (QE × perturbation × *W*_MF_ × *t*_MF_: *b* = –2.21×10^−1^, *s.e.m.* = 4.97×10^−2^, *p* < 0.0001). However, QE also increased error in combination with perturbation and *t*_MF_ even in the absence of any moderating effect of *W*_MF_ (QE × perturbation × *t*_MF_: *b* = 8.62×10^−3^, *s.e.m.* = 4.18×10^−3^, *p* < 0.05). This result supports Hypothesis-1a.

Despite the absence of a simple QE × perturbation interaction, the eye height perturbation produced a greater model-predicted increase in radial error for QE-trained participants. To illustrate this point, we can compare the model predictions for participants in each condition with high and low *W*_MF_ and *t*_MF_ (with ‘low’ and ‘high’ values defined as 1^st^ and 3^rd^ quartiles). For participants with ‘low’ *W*_MF_, the eye height manipulation prompted 2.27% to 11.14% increase in radial error in QE-trained compared to technical-trained participants for low to high *t*_MF_, respectively. For participants with ‘high’ *W*_MF_, the eye height manipulation prompted 8.80% to 18.23% increase in radial error in QE-trained compared to technical-trained participants for low to high *t*_MF_, respectively.

### 3.1.4. Testing Hypothesis-1b: the QE group with greater multifractal nonlinearity in sway showed a greater initial but progressively slower increase in radial error due to the perturbation of eye height

The interactions of QE × trial_block_ with multifractal nonlinearity followed a similar but weaker pattern as the interactions of QE × trial_block_ with perturbation. Much like the perturbation of eye height, the interaction of QE with *t*_MF_ accentuated the radial error late in a block of ten trials (QE × trial_block_ × *t*_MF_): it weakened the linear term (*b* = –3.73×10^−2^, *s.e.m.* = 1.46×10^−2^, *p* < 0.05), contributing to a negative-quadratic peak in the middle of the block (*b* = –1.85×10^−1^, *s.e.m.* = 1.66×10^−2^, *p* < 0.0001) and a positive cubic increase at the end of the block (*b* = 1.13×10^−1^, *s.e.m.* = 1.73×10^−2^, *p* < 0.0001). Hence, *t*_MF_ and perturbation showed independent effects on the progression of error across trials in block.

However, the similarity of these effects independently on error across trials within a block did not extend to their interaction. Indeed, most relevant to testing Hypothesis-1b is that the interaction of perturbation and *t*_MF_ served to speed up the growth of error, and QE slowed it down across the block. Coefficients for perturbation × trial_block_ × *t*_MF_ indicated that in the face of perturbation, multifractal nonlinearity removed the observed peak in error due to earlier negative quadratic effects (*b* = 3.83×10^−1^, *s.e.m.* = 5.28×10^−2^, *p* < 0.0001) and increased error across the linear (*b* = 4.07×10^−1^, *s.e.m.* = 5.20×10^−2^, *p* < 0.0001)and cubic terms (*b* = 5.39×10^−1^, *s.e.m.* = 4.45×10^−2^, *p* < 0.0001). On the other hand, coefficients for perturbation × trial_block_ × *t*_MF_ indicated that, in the face of perturbation for the QE participants, the linear increase was shallower (*b* = –2.71×10^−1^, *s.e.m.* = 6.11×10^−2^, *p* < 0.0001), the positive quadratic growth further removed the observed peak in error (*b* = 4.24×10^−1^, *s.e.m.* = 6.14×10^−2^, *p* < 0.0001), and the cubic term was negative (*b* = –7.90×10^−1^, *s.e.m.* = 5.79×10^−2^, *p* < 0.0001).

The outcome of these differences distinguishes how multifractal nonlinearity predicted changes in adaptive responses to the perturbation of eye height. Greater *t*_MF_ exhibited less adaptive response to the perturbation in the QE case (i.e., initially greater error and a smaller change in error across trials), and less-but-still-nonzero *t*_MF_ predicted a more gradual appearance of error and then a subsequent decay of this error (Fig. 5*b*), suggesting better adaptation. It is evident in figure 5*b* that although the error in the high-*t*_MF_ cases with the perturbation was smaller than the delayed increase in the low-*t*_MF_ cases with the perturbation, the high-*t*_MF_ cases showed more sustained error and no adaptation to the perturbation relative to high-*t*_MF_ cases without the perturbation. This result supports Hypothesis-1b.

We include example plots of model predictions for individual participants (Fig. 6).

**Figure 6.**
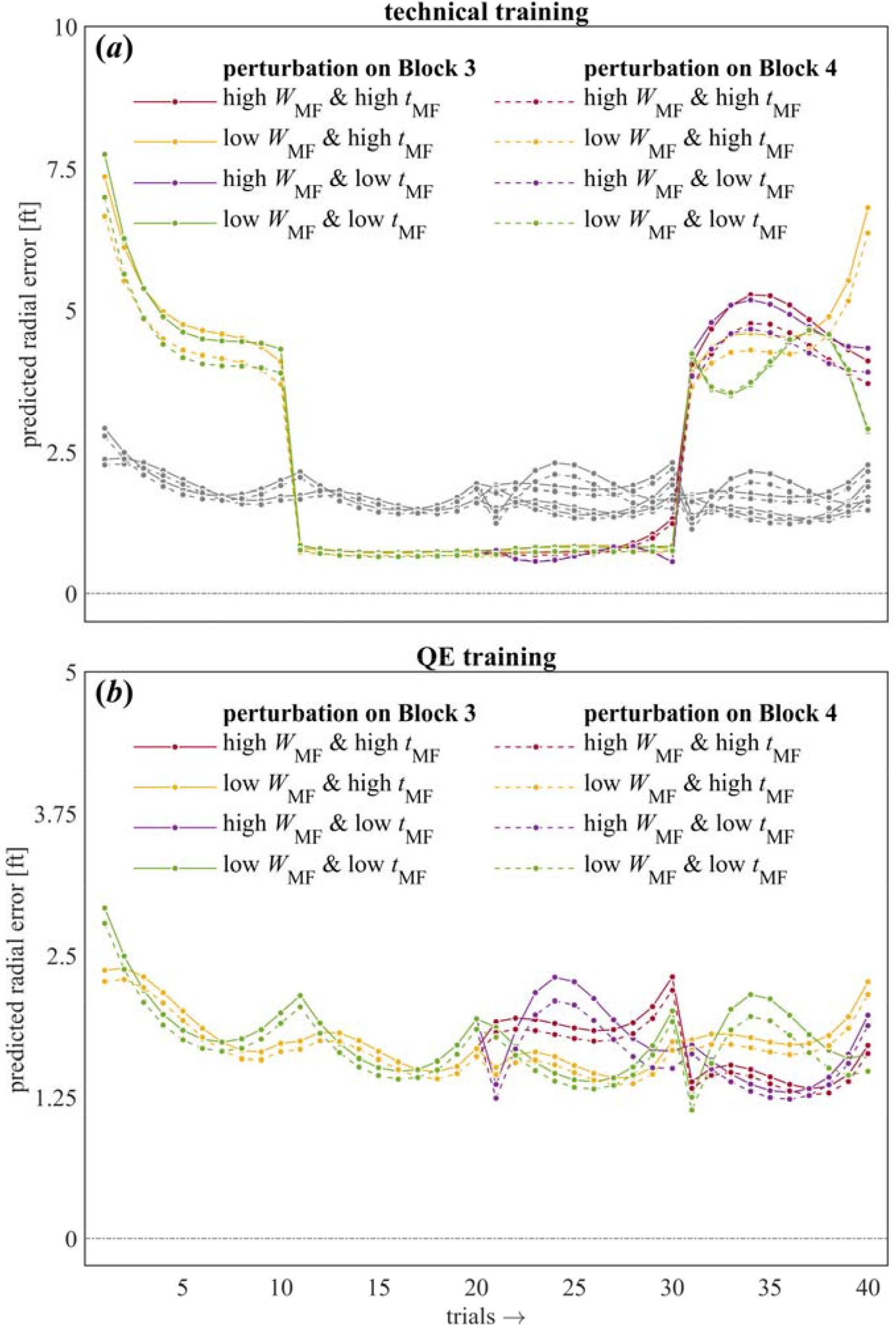
The observed trial-by-trial radial error and trial-by-trial model predictions on the same axis for four participants each from the technical (top four panels) and QE-trained (bottom four panels) groups, respectively. The representation includes two participants each with the highest (left panels) and lowest (right panels) Pearson’s correlation coefficient *r* indicating the strength of relationship between the observed and predicted radial errors.

### 3.2. Logistic regression of ‘makes’ vs. misses showed less sensitivity to effects

Poisson regression may be too ornate for some readers. However, in contrast to Poisson modeling of a continuous outcome measure of performance, the investment of logistic regression to model dichotomous variable limits the number of valid predictors (e.g., van Smeden et al., 2016) and so affords us less statistical power with which to test for changes in the performance. A mixed-effect logistic regression model of successful putting (i.e., the ‘makes’) returned the coefficients reported in Table 3. Predictors for *t*_MF_ and for interactions of QE with all terms other than perturbation failed to improve model fit significantly and so were omitted from the final model. This model failed to return an effect of QE (*b* = 0.20, *s.e.m.* = 0.30, *p* = 0.51), of the perturbation alone (*b* = –0.22, *s.e.m.* = 0.43, *p* = 0.60) or the interaction of QE × perturbation (*b* = –1.16, *s.e.m.* = 0.65, *p* = 0.07).

**Table 3.** Logistic regression modeling of successful shots.

| **predictor** | ***b*** | ***s.e.m.*** | ***p*<sup>\*</sup>** |
| --- | --- | --- | --- |
| intercept | −2.85 | 0.48 | < **0.0001** |
| QE | 0.20 | 0.30 | 0.51 |
| perturbation | −0.22 | 0.43 | 0.60 |
| QE × perturbation | −1.16 | 0.65 | 0.07 |
| block | 0.24 | 0.13 | **0.06** |
| *RMS*<sub>Acc</sub> | 4.05 | 1.34 | < **0.01** |
| *W*<sub>MF</sub> | −7.25 | 2.94 | < **0.05** |

\*boldfaced values indicate statistical significance at the alpha level of 0.05.

We found effects of experience and trial-by-trial sway. Block showed a positive effect (*b* = 0.24, *s.e.m.* = 0.13, *p* = 0.06), suggesting that odds of a successful putt was 1.27 times higher with each successive block of trials. *RMS*_acc_ showed a positive effect (*b* = 4.05, *s.e.m.* = 1.34, *p* < 0.01), and total multifractality *W*_MF_ showed a negative effect (*b* = –7.25, *s.e.m.* = 2.94, *p* < 0.05), suggesting that successful putts were more likely with more variable and less multifractal accelerations of sway. These effects resembled some of those found in the Poisson regression model, but it appears that the continuous measure of radial error afforded more sensitivity within which to model significant fixed effects.

## 4. Discussion

We tested the effects of postural stability and its endogenous multifractal fluctuations on a continuous outcome of putting performance (radial error) with and without QE training. We hypothesized that QE-trained participants’ response to an eye height perturbation would depend on the multifractality attributable to nonlinear interactions across scales, *t*_MF_. That is, more multifractal fluctuations in postural sway would make eye height perturbation a greater threat to QE-trained performance but not to technically trained performance. Specifically, we predicted that lower estimates of *t*_MF_ would coincide with a more adaptive response, i.e., with slower growth of radial error and subsequent decay of radial error over trials. Results supported this hypothesis: when we perturbed eye height, accuracy with QE was associated with reducing multifractal nonlinearity in postural sway. This model included a negative main effect of QE training, indicating more accurate putts in radial error. However, despite the absence of a simple interaction of QE with clogs, QE’s contingency on multifractal aspects of postural sway entailed a greater decrement in accuracy for QE-trained participants following eye height perturbation.

A limitation of these results is that our focus on the postural support of QE-trained performance came at the expense of focusing on gaze tracking. We used QE training as a manipulation, but we failed to include a manipulation check. The absence here of gaze-tracking data leaves us unable to explain with complete confidence why a logistic model of putting performance construed dichotomously (e.g., as ‘makes’ vs. ‘misses’) failed to find a significant effect of QE. On the one hand, some participants may have followed QE instruction incompletely. On the other hand, a good reason why the main effect may not appear is the brevity of our design: whereas recent analogues of our research design already in the literature required participants to conduct 4000 putts over 7 days (Moore et al., 2012), we only sampled 40 putts in one hour-long session. Radial error may be sufficiently sensitive to show QE’s reliable effect in a more compact design than dichotomous measures. Future research would no doubt benefit from incorporating gaze-tracking into similarly compact experimental designs.

These results offer two major insights for research into perception-action in visual aiming tasks. First, it shows that QE training has roots in postural sway, vindicating and elaborating earlier proposals that quiet eye is not simply about stabilizing the eyes (Liphart et al., 2015; Williams, 2016). Indeed, it has long been known that simply stabilizing the retinal position compromises the persistence of a visual image (Barlow, 1963; Coppola & Purves, 1996; Ditchburn & Ginsborg, 1952). And contrary to a colloquial understanding of ‘quiet eye,’ more recent work confirms that those small movements within gaze fixations called ‘microsaccades’ might prevent fading (Martinez-Conde et al., 2006) and support visual attentional processes (Krueger et al., 2019). We might draw a comparison between the ‘quiet’ suggested by ‘quiet eye’ and that indicated by ‘quiet standing.’ To be precise, we mean to draw the comparison of ‘quiet eye’ specifically only to ‘quiet standing’ in the case of instructing participants to ‘stand as still as possible’ (i.e., rather than to ‘stand as relaxed as possible’; (Houdijk et al., 2015). Both ‘quietudes’ provide the needed instruction to a participant or athlete to ‘please move as little as possible,’ but below the polite clarity for verbal instruction, both ‘quietudes’ depend upon a rich texture of fluctuations. Throughout the body, fluctuations support exploration during the ‘still as possible’ quiet standing (I.-C. Lee et al., 2019; Murnaghan et al., 2016; Palatinus et al., 2013, 2014).

Here, the theoretical ground may shift underfoot as we try to see a path through these considerations. Do we mean to say that the achievement of a precision demand could be exploratory? At first blush, it could sound absurd that something not moving is also exploring. However, here, we may hit upon a deeper divergence between ecological vs. information-processing perspectives on QE. To the point, it is true that fixing the gaze and standing as still as possible are themselves accomplishments of motor control. However, this achievement does not entail that either quiet eye or quiet stance are the results. Unless we are much mistaken, it is the external focus and the intention of getting the ball into the net or hole or target that warrants quiet eye in the first place. That is to say, the quiet eye is a means to an end, and the end is never just quiet eye.

Precision might invite the appearance of achieving fixity, but achieving fixity might be no guarantee against exploration. For a contrary example, ‘static holding’ is a haptic case of holding an object ‘as still as possible,’ and it is widely recognized as a valid exploratory procedure for haptic perception (Burton & Turvey, 1990; Carello et al., 1992; Lederman et al., 1996). The only debates are what kind of information this apparent-fixity mode of exploration has access to. Crucially, apparent fixity is not necessarily actual fixity, and while there is no need for the body to explore the quiet-eye achievement of ocular fixity, it would strike us as a strange eventuality to rule quiet-eye strategy out as a strategy pertaining to exploration. Quiet eye resembles to us an ocular version of static holding that holds the visual image as close as possible. As noted above, despite their apparent fixity, fixations are a class of eye movements, and these fixational eye movements are exactly what the quiet-eye strategy uses to build a motor program. Certainly, if the QE strategy is collecting information that supports the construction of a motor program, ruling quiet eye out as a form of exploration would strike us as a problematic move. For instance, consider the pioneering astronomers gazing up at the heavens, aiming a telescope as quietly as possible at a star or planet. Can we tell them that, no, because they are engaged in precision work, they are not exploring? We suspect not. Again, to summarize this point: a precision demand can change motor behavior (Balasubramaniam et al., 2000), but that change is not itself the end of the story. Despite experimenter intentions to train it, quiet eye is not a goal in itself but only a strategy for informing the participant’s accurate behavior beyond their bodies for precision wherever they direct their external focus.

Of course, the preceding proposal that QE is exploratory remains conjecture that reaches beyond the current evidence. Demonstrating an exploratory structure in QE will need empirical evidence from future research to back up the preceding metaphor and analogy. Specifically, empirical validation would require at least one of the following: (1) eye movement data indicating exploratory movements within the fixation time that are similar or relational to those from the haptic or postural research, (2) performance outcome data that indicates information is obtained that is of value during the quiet eye fixation period, and/or (3) self-reporting of the subjects (perhaps quantified through perceptual judgments) indicating information gained following QE training that differs from those who do not go through the training. We look forward to future research to the opportunity to test these ideas and revisit the question of whether QE is exploratory behavior.

We may summarize this first insight as follows: The value of QE is not necessarily a net reduction of fluctuations but rather a way to train athletes to orient their movements towards the optic flow generated by postural sway. Whereas the postural-kinematic hypothesis had previously only referred to longer movement duration (Gallicchio & Ring, 2020), we now report that movements are not merely slower but evince a sort of nonlinearity constricting the degrees of freedom across scales of the movement-system hierarchy. In one of the subsections below, we discuss how this orienting towards optic flow might coordinate with mechanisms invoked by the more elaborate visual hypothesis (e.g., Gallicchio et al., 2016, 2017).

The second insight from the present results is that the multifractality in sway is an essential aspect of postural adaptations contributing to visually-guided action. It was already known that damping out postural perturbations could occasion a ratcheting down of multifractality in postural sway (Kelty-Stephen, 2018). The present results indicate that this reduction of multifractality is associated with more accurate responses in visually-guided aiming. Note that this reduced multifractality remains different from corresponding surrogates. Hence, nonlinear interactions across scales can constrict movement variability, making reduced multifractality more adaptive. This distinction is a departure from any shorthand presumption that ‘more multifractal is better’ supported by early studies of heart rate variability (HRV; Ivanov et al., 2001). Indeed, this departure reflects a growing clarity about the value of multifractal fluctuations for predicting and explaining outcomes in perception-action.

### 4.1. Clarifying the value of multifractal fluctuations for perceiving-acting systems

The shorthand ‘more multifractal is better’ presumptions from early studies on HRV (Ivanov et al., 2001) have always underestimated the more elaborate systematicity in the physiology of the human movement system with upright posture. Early work proposed that a ‘loss of [fractal or multifractal] complexity’ would coincide with aging or with pathology (Lipsitz & Goldberger, 1992). Indeed, early evidence showed that less healthy outcomes co-occurred with more-than-fractal levels of temporal correlation (Lipsitz, 2002). However, the task has always been known to influence multifractality in upright posture: multifractality of quiet standing can reduce with pathology (Morales & Kolaczyk, 2002; Shimizu et al., 2002), but multifractality in gait can also increase with pathology (Muñoz-Diosdado, 2005), as well as with walking speeds faster or slower than self-selected comfortable speed (West & Scafetta, 2003). Over the lifespan, the maturation of gait involves loss of multifractality, beginning with the stabilization of toddler gait into young adult gait and continuing into older age (Ashkenazy et al., 2002). And this lifelong progression of multifractality in gait actually is at odds with more recent evidence that multifractality in HRV remains stable with healthy aging (Schmitt & Ivanov, 2007).

Militating strongly against simple ‘more multifractal is better’ kind of presumptions are a variety of distinctions. Some of these distinctions may sound patently obvious but need recognition if we want to identify for what, after all, multifractality is good. These distinctions include, for instance, that hearts are not upright bipedal bodies, that age is not itself a pathology, and that walking is not standing. The remaining distinctions are more subtle. In the promise of fractal/multifractal analysis, the discourse about ‘complexity,’ ‘dynamical stability,’ and the role of fractality/multifractality in ‘optimality’ (Friston et al., 2012; Oftadeh et al., 2014; Schlesinger & Klafter, 1986) has made no claims on all definitions of optimality and so has not been responsible for acknowledging that some multifractality is not optimal (Reynolds, 2015). Indeed, studies in behavioral sciences have always justified the capacity of fractal variety to offer new insights (Diniz et al., 2011; Dixon et al., 2012). Any notion of a privileged level or direction of fractal variety (Kello, 2018) might be best understood as a reaction to critical, unreceptive perspectives resistant to seeing any fractality or multifractality appear in behavioral-science discourse at all (Wagenmakers et al., 2012). Measurements may wax and wane in their multifractal structure. The only case in which we think more multifractality is always better than less is in its inclusion as a construct in the scientific discourse on the coordination of perceiving-acting systems.

### 4.2. The value of multifractal fluctuation is specific to task constraints

What is coming into focus is that the value of nonlinearity-driven multifractality depends on task context. More multifractal nonlinearity is adaptive when tasks invite exploration and anticipation. Less multifractal nonlinearity is adaptive when the task involves constraining behavior, drawing degrees of freedom inward around a base of support. As an example of the former, accumulating information as a reader can require keeping an open mind to follow the discourse wherever it may lead, and more multifractality *t*_MF_ in the pacing of word-by-word reading supports more fluent reading if the story has a twist in its plot (Booth et al., 2018). Similarly, more multifractality *W*_MF_ in circle-tracing behaviors make for poor precision of tracing but is associated with better performance on neurophysiological tests of flexibility with rule switching (Kelty-Stephen et al., 2016). The freewheeling benefits of more multifractality can also compete alongside the constraint-, repetition-promoting benefit of less multifractality. In a Fitts task, greater multifractality *t*_MF_ in hand and head movements predicted less stable contact with the targets, but it became a predictor of more stable contact when participants had the diffuse warning that they might be asked to close their eyes and continue the task (Bell et al., 2019). Hence, what would have upset task performance became a resource for exploring the task space for the eventuality of a major loss of sensory information. And now, we see that a less multifractal postural system can be more effective (e.g., smaller *t*_MF_ in Kelty-Stephen (2018); and smaller *W*_MF_ in Koslucher et al. (2016) and Li et al. (2020)) and, as present findings indicate, more accurate in how it orients behaviors to a visually-guided aiming task.

Some of the task-specificity remains unclear, e.g., manual wielding of an object is more accurate with greater multifractality in postural sway, but hefting an object to perceive heaviness compared to a reference object is more accurate with less multifractality (Mangalam & Kelty-Stephen, 2020). This difference could have to do with the qualities of length and heaviness, but the simpler interpretation could be that intending to perceive length directs attention away from the center of pressure and intending to perceive heaviness directs attention inwards toward the center of pressure, lending length perceptions and heaviness perception to less and more postural constraint, respectively. This issue of directing attention is again a reason we will need to consider how to link this postural-kinematic work on QE back with the visual-hypothesis work on QE.

The fact that reduced multifractality can be beneficial offers a unique distinction between possibly two related but distinct interpretations of multifractal results: one as a biomarker of pathology and the other as an adaptation. Indeed, the loss of multifractality can be a biomarker for diagnosing a pathology (Meyer et al., 2003). The physiological wisdom informing this usage is that the heart has to be, in effect, poised to absorb and rebound from perturbations, avoiding fragility due to insult at any single characteristic scale (e.g., Lipsitz & Goldberger, 1992). Implicit in this wisdom is that, within the healthy body, the heart’s task is deeply anticipatory and exploratory, much like those task settings in which whole organisms benefit from more multifractality. However, concluding that loss of multifractality is strictly symptomatic of postural systems with deficits (Shimizu et al., 2002) may be premature. What looks like a deficit may sometimes be a creative constraint, built and used to solve a task problem (Harrison & Stergiou, 2015; Holt et al., 2010; Vaz et al., 2017). The present results indicate that reduced multifractality could be an adaptive response preserving stability and promoting task performance in the face of abrupt change in the organism’s mechanical relationship with the base of support. Hence, far from being the signature of diseased posture, reductions in multifractality that maintain a significant *t*_MF_ may be an advantageous loss of multifractality. Though more commonly known for their capacity to amplify variability, nonlinear interactions across scales acting context-sensitively may also restrain motoric degrees of freedom so as to achieve the task goal.

Different task constraints prompt different modes of movement variability, and if multifractal depictions of the body seem to physicalize or mechanize the movement system, they do so only in the way that nonlinear-dynamical physical mechanisms might better explain context-sensitive behavior. Surely, good-faith pursuit into nonlinear-dynamical complexity should avoid any catch-all simplification (e.g., ‘more is better, less is worse’) and keep a critical eye on the reference frames of nonlinear-dynamical results. Nonlinear dynamics has long profited from considering its measures in different reference frames: experimentally breaking apart the coincidence of relative-phase in spatial frames from relative-phase in muscular frames revealed novel, more generic structure in the Haken-Kelso-Bunz (1985) law for multi-limb coordination (Park & Turvey, 2008). Acknolwedging the task-based reference frames for our interpretation of multifractal estimates may be equally crucial for generalizing predictions for a multifractal foundation for perception-action. Low multifractality in the heart signals a bad prognosis to the clinician when the task is cardiovascular circulation, but low multifractality in posture supports good task performance in visually-guided aiming, particularly for an organism beset by malevolent experimenters uprooting their optic flow.

### 4.3. The value of multifractality is also specific to different tissues of the body

Just as different tissues of the same body can differently support the same task, we expect that the multifractal structure spanning these different tissues supports overarching integrity to the organismal-level behavior differently (Stoffregen et al., 2017). Here, comparing present results with similar work using a relatively less-mechanical, visual perturbation (Carver et al., 2017) may tee up considerations for future work that could elaborate an explanation of QE by integrating the postural-kinematic hypothesis with the visual hypothesis. The present results resemble those of Carver et al. (2017) who found that, without any training, more multifractality in head sway led to greater error in a low-difficulty Fitts task (compare *W*_MF_ in Table 2) but led to smaller error in the case of a visual perturbation (compare perturbation × *W*_MF_ in Table 2). So, multifractality in head sway became a stabilizing, accuracy-promoting asset to task performance when the target was far out of reach. Future research could probe the visual hypothesis for QE for multifractal foundations, and based on Carver et al.’s (2017) results, we predict that QE-trained performance would improve with greater multifractality in head sway or eye movements. Monofractal analysis of eye movements has successfully predicted visual attention to text (Wallot et al., 2015), and multifractality of gaze fluctuations may support an extension of the multifractal support for QE. The present results do not include gaze data, but we hope it is informative, especially in support of the postural-kinematic hypothesis, that the QE instructions enlist the nonlinear interactions across scales of the movement system without drawing the athlete’s attention to the body.

Ultimately, this task- and anatomically-specific portrait of multifractal fluctuations rests on two even more generic facts. First, multifractal fluctuations spread across the body, and second, estimates of this spread predict the accuracy of the perceptual outcomes (Mangalam, Carver, et al., 2020a, 2020b). Hence, rather than inventorying body parts and task constraints prompting different values of multifractality, the longer view of investigating visually-guided aiming must examine the flow of multifractal fluctuations. Carver et al. (2017) found that multifractality spread from hip to head, as well as from head and hip toward the throwing hand. Indeed, in all these studies, the body appears to direct multifractal fluctuations towards those body parts most clearly engaged in the focal tasks. We might expect similar results in golf putting. QE training may promote this spread by reducing intentional movements of the head and eyes that might interfere with upstream flows of multifractality. The direction of attention towards a distant target in the visual field may depend on stabilizing the postural grasp of the surface underfoot and, perhaps, shunting multifractal fluctuations toward the head and eyes. Indeed, executive functions directing attention to specific features of the visual field benefits from a rich substrate of multifractal fluctuations in the head (Kelty-Stephen & Dixon, 2014), hand (Anastas et al., 2014; Stephen et al., 2012), and brain (Kardan, Adam, et al., 2020; Kardan, Layden, et al., 2020) that vary systematically with access to and mastery of the rules supporting task performance. When we consider that eye movements show multifractality (Kelty-Stephen & Mirman, 2013) and that variability in the monofractal structure of eye movements can predict visual attention (Wallot et al., 2015), we see early glimpses of a multifractal bridge from the postural-kinematic support to the visual support for QE.

The novelty of multifractal modeling to behavioral sciences can make this promise of a multifractal infrastructure supporting visually guided behaviors sound too ethereal and misty to be practical. But we can root this proposal in the known physiology spanning the very same bones, muscles, bones, joints, brain, and eyes whose coordination generates the winning golf putt. Specifically, the connective tissues composing the fascia operate to coordinate context-sensitive behavior across so wide a variety of scales (e.g. (Ingber, 2006)) that multifractality is one of the few current compelling frameworks for modeling how it might support perception-action (Schleip et al., 2014; Turvey & Fonseca, 2014). The flow of multifractal fluctuations through the body could inform future explanations of how QE training and other visual aiming tasks depend on this bodywide network of connective tissues.

## Supplementary materials

**Supplementary Table S1.** Subset of Poisson regression model output including main and interaction effects of time and perturbation of eye height.

