## Supplementary Table 1 for "Multifractality in postural sway supports quiet eye training in aiming tasks: A study of golf putting"

**Supplementary Table S1.** Subset of Poisson regression model output including main and interaction effects of time and perturbation of eye height.

| predictor | *b* | *s.e.m.* | *p*[*] |
|---|---|---|---|
| intercept | $9.04\times10^{0}$ | $3.29\times10^{-1}$ | **< 0.0001** |
| $\text{trial}_{exp}$(Linear) | $4.70\times10^{1}$ | $2.26\times10^{0}$ | **< 0.0001** |
| $\text{trial}_{exp}$(Quadratic) | $3.08\times10^{1}$ | $1.20\times10^{0}$ | **< 0.0001** |
| $\text{trial}_{exp}$(Cubic) | $-1.78\times10^{0}$ | $1.54\times10^{-1}$ | **< 0.0001** |
| $\text{trial}_{block}$(Linear) | $2.17\times10^{1}$ | $9.34\times10^{-1}$ | **< 0.0001** |
| $\text{trial}_{block}$(Quadratic) | $9.60\times10^{-1}$ | $2.63\times10^{-1}$ | **< 0.001** |
| $\text{trial}_{block}$(Cubic) | $-1.62\times10^{0}$ | $2.57\times10^{-1}$ | **< 0.0001** |
| block | $-1.41\times10^{0}$ | $1.34\times10^{-1}$ | **< 0.0001** |
| block × $\text{trial}_{block}$(Linear) | $-1.36\times10^{1}$ | $5.83\times10^{-1}$ | **< 0.0001** |
| block × $\text{trial}_{block}$(Quadratic) | $-1.10\times10^{0}$ | $1.10\times10^{-1}$ | **< 0.001** |
| block × $\text{trial}_{block}$(Cubic) | $6.40\times10^{-1}$ | $1.08\times10^{-1}$ | **< 0.0001** |
| QE | $-4.40\times10^{0}$ | $4.81\times10^{-1}$ | **< 0.0001** |
| QE × $\text{trial}_{exp}$(Linear) | $-6.07\times10^{1}$ | $3.54\times10^{0}$ | **< 0.0001** |
| QE × $\text{trial}_{exp}$(Quadratic) | $-3.21\times10^{1}$ | $1.86\times10^{0}$ | **< 0.0001** |
| QE × $\text{trial}_{exp}$(Cubic) | $1.03\times10^{0}$ | $2.18\times10^{-1}$ | **< 0.0001** |
| QE × $\text{trial}_{block}$(Linear) | $-2.41\times10^{1}$ | $1.41\times10^{0}$ | **< 0.0001** |
| QE × $\text{trial}_{block}$(Quadratic) | $9.35\times10^{-1}$ | $3.79\times10^{-1}$ | **< 0.05** |
| QE × $\text{trial}_{block}$(Cubic) | $2.24\times10^{0}$ | $3.54\times10^{-1}$ | **< 0.0001** |
| QE × block | $1.72\times10^{0}$ | $1.96\times10^{-1}$ | **< 0.0001** |
| QE × block × $\text{trial}_{block}$(Linear) | $1.56\times10^{1}$ | $8.99\times10^{-1}$ | **< 0.0001** |
| QE × block × $\text{trial}_{block}$(Quadratic) | $1.31\times10^{0}$ | $1.63\times10^{-1}$ | **< 0.0001** |
| QE × block × $\text{trial}_{block}$(Cubic) | $-5.57\times10^{-1}$ | $1.54\times10^{-1}$ | **< 0.001** |
| Perturbation | $3.90\times10^{-2}$ | $3.01\times10^{-1}$ | 0.90 |
| QE × perturbation | $8.73\times10^{-2}$ | $4.26\times10^{-1}$ | 0.84 |
| perturbation × $\text{trial}_{block}$(Linear) | $1.58\times10^{0}$ | $3.02\times10^{-1}$ | **< 0.0001** |
| perturbation × $\text{trial}_{block}$(Quadratic) | $1.25\times10^{0}$ | $3.08\times10^{-1}$ | **< 0.001** |
| perturbation × $\text{trial}_{block}$(Cubic) | $-2.42\times10^{0}$ | $2.97\times10^{-1}$ | **< 0.0001** |
| QE × perturbation × $\text{trial}_{block}$(Linear) | $-1.16\times10^{0}$ | $4.44\times10^{-1}$ | **< 0.01** |
| QE × perturbation × $\text{trial}_{block}$(Quadratic) | $-5.71\times10^{0}$ | $4.43\times10^{-1}$ | **< 0.0001** |
| QE × perturbation × $\text{trial}_{block}$(Cubic) | $3.39\times10^{0}$ | $4.31\times10^{-1}$ | **< 0.0001** |

[*]boldfaced values indicate statistical significance at the alpha level of 0.05.
